# cgRNASP: coarse-grained statistical potentials with residue separation for RNA structure evaluation

**DOI:** 10.1101/2022.03.13.484152

**Authors:** Ya-Lan Tan, Xunxun Wang, Shixiong Yu, Bengong Zhang, Zhi-Jie Tan

## Abstract

Knowledge-based statistical potentials are very important for RNA 3-dimensional (3D) structure prediction and evaluation. In recent years, various coarse-grained (CG) and all-atom models have been developed for predicting RNA 3D structures, while there is still lack of reliable CG statistical potentials not only for CG structure evaluation but also for all-atom structure evaluation at high efficiency. In this work, we have developed a series of residue-separation-based CG statistical potentials at different CG levels for RNA 3D structure evaluation, namely cgRNASP, which is composed of long-ranged and short-ranged interactions by residue separation. Compared with the newly developed all-atom rsRNASP, the short-ranged interaction in cgRNASP was involved more subtly and completely through explicitly adding the interactions between nearest neighbor residues and between next-nearest ones. Our examinations show that, the performance of cgRNASP varies with CG levels and compared with rsRNASP, cgRNASP can have similarly good performance for extensive test datasets and slightly better performance for the realistic RNA-Puzzles dataset. Furthermore, cgRNASP is strikingly more efficient than all-atom potentials such as rsRNASP, and can be apparently superior to other all-atom statistical potentials and scoring functions trained from neural networks for the RNA-Puzzles dataset. cgRNASP is available at https://github.com/Tan-group/cgRNASP.

## Introduction

RNAs have critical biological functions such as gene regulations and catalysis (1,2), and their functions are generally coupled to their structures (3,4). Consequently, the knowledge of RNA structures, especially 3-dimensional (3D) structures, is crucial for understanding RNA biological functions (5,6). Due to the huge cost of experimental measurements, high-resolution 3D structures of RNAs deposited in protein data bank (PDB) database are still limited (7). Complementary to experiments, various computational models have been developed to predict RNA 3D structures *in silico* (8,9), and these models can be roughly classified into fragment-assembly-based ones and physics-based ones (10-46). A computational model for 3D structure prediction generally requires a reliable energy function for evaluating/assessing predicted structure candidates (47,48). Therefore, a reliable energy function or a reliable structure evaluation is very important for a computational model in RNA 3D structure prediction.

For proteins, knowledge-based statistical potentials derived from experimental structures deposited in the PDB database have been shown to be rather efficient and effective for the quality evaluation/assessment of protein 3D structures, protein-protein and protein-ligand docking (49-67). For RNA 3D structure evaluation, several statistical potentials have been developed based on different reference states (68-72). Bernauer et al have derived differentiable statistical potentials (KB) at both all-atom and coarse-grained levels based on the quasi-chemical approximation reference state (68). Capriotti et al have built all-atom and coarse-grained statistical potentials (RASP) based on the averaging reference state (69). Wang et al have derived a combined distance- and torsion angle-dependent all-atom statistical potential (3dRNAscore) based on the averaging reference state (70). Zhang et al have proposed an all-atom statistical potential based on the finite-ideal-gas reference state (DFIRE-RNA) (71). Recently, we have made a comprehensive survey on the six existing reference states widely used for proteins through building six statistical potentials based on the same training dataset, and found that the finite-ideal-gas and random-walk-chain reference states are modestly better than other ones in identifying native structures and ranking decoy structures (72). However, the existing traditional statistical potentials only achieve a poor performance for the realistic test dataset — RNA-Puzzles dataset (73,74). Beyond traditional statistical potentials, machine/deep learning approaches have been employed for RNA 3D structure evaluation (75,76). Despite of “black-box” training/learning process, RNA3DCNN built through 3D convolutional neural networks exhibits remarkably improved performance in identifying native structures for RNA-Puzzles dataset compared with the top traditional statistical potentials (75), and the newly developed scoring function of ARES from deep neural network based on training data from FARFAR2 showed rather good performance for evaluating structures from FARFAR2 (76). Very recently, we have developed an all-atom statistical potential of rsRNASP by distinguishing short- and long-ranged interactions at residue separation level, and for the test datasets from various 3D structure prediction models, rsRNASP has visibly improved performance than existing statistical potentials and scoring functions from neural networks (47,77).

Very importantly, for reducing conformational space and improving computational efficiency, almost all existing physics-based models for RNA 3D structure prediction are based on different-level CG representations rather than the all-atom representation, including SimRNA (43,44), iFold (29,30), NAST (31), IsRNA (26,27), Vfold (23,25), HiRe-RNA (40,41), oxRNA (42), RACER (45,46), and our CG model with salt effect (33,37). Consequently, a reliable CG statistical potential is crucially important for a CG-based 3D structure prediction model rather than an all-atom-based one. However, existing CG statistical potentials (e.g., CG KB (68) and CG RASP (69)) are still at very low performance and recently developed statistical potentials/scoring functions with relatively good performance (e.g., rsRNASP (77), RNA3DCNN (75), ARES (76) and DFIRE-RNA (71)) are all based on the all-atom representation. Therefore, until now, reliable CG statistical potentials are still highly required at different CG levels. Such CG statistical potentials can be very useful not only for CG structure evaluation but also for all-atom structure evaluation at high efficiency.

In this work, we have developed a series of distance-dependent CG statistical potentials based on residue separation for RNA 3D structure evaluation at different CG levels, named cgRNASP, which is composed of short- and long-ranged potentials distinguished by residue separation. In cgRNASP, beyond the newly developed all-atom rsRNASP, the interactions between nearest neighbor residues and between next-nearest ones were explicitly added in the short-ranged interaction. The performance of cgRNASP can have slightly better performance than the all-atom rsRNASP for the realistic RNA-Puzzles dataset and strikingly higher evaluation efficiency than all-atom statistical potentials such as rsRNASP. Moreover, cgRNASP is apparently superior to other all-atom traditional statistical potentials and scoring functions trained from neural networks for the RNA-Puzzles dataset.

## Materials and methods

### Coarse-grained representations

First, a survey was made on existing physics-based models for RNA 3D structure prediction to figure out which heavy atoms were used for representing CG atoms in these models, and for convenience, we only considered real heavy atoms rather than dummy atoms since some CG-based models involved the (mass) centers of certain atom groups as dummy CG atoms. As shown in Table 1, P and C4’ atoms were used most widely to describe RNA backbones, and N9 atom for purine or N1 atom for pyrimidine was used frequently to (partially) describe bases. According to Table 1, we developed our residue-separation-based CG statistical potentials (cgRNASP) at several CG levels: (i) three CG beads at P, C4’, and N9 atoms for purine (or N1 atom for pyrimidine); (ii) two CG beads at P and C4’ atoms, and (iii) one CG bead on C4’ atom, which were also illustrated in Fig. 1. Correspondingly, for simplicity, our CG statistical potentials are named as cgRNASP, cgRNASP-PC, and cgRNASP-C, respectively. The first potential was named in such way since the potential of 3-bead representation was regarded as a representative one of the CG potentials.

**Table 1.** Heavy atoms for CG beads used in cgRNASP and in existing CG-based RNA 3D structure models

| Heavy atoms for CG beads | No of CG beads | CG-based structure prediction models |
| --- | --- | --- |
| P, C4', N1 (or N9) <sup>a</sup> | 3 | SimRNA [43], Vfold [21, 23], Shapiro's model [78]; A CG model with salt effect [32, 33] |
| P, C4' | 2 | SimRNA [43], Vfold [21,23], Shapiro's model [78]; HiRE-RNA [39,40], IsRNA [25,26], RACER [41,44], A k-state virtual bond model [79], A CG model with salt effect [32,33] |
| C4' | 1 | SimRNA [43], Vfold [21,23], Shapiro's model [78]; HiRE-RNA [39,40], IsRNA [25,26], RACER [41,44], A k-state virtual bond model [79], A CG model with salt effect [32,33] |

<sup>a</sup> P, C4', and N1 are for pyrimidine, and P, C4', and N9 are for purine.

**Figure 1.**
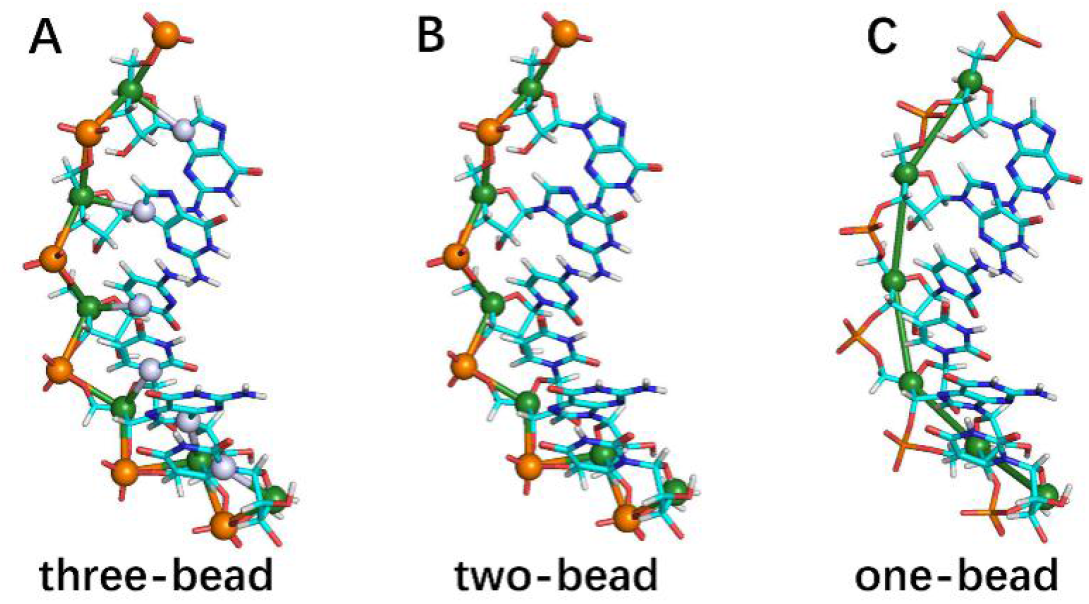
Illustration for different coarse-grained (CG) representations used in developing cgRNASP for RNA 3D structure evaluation. (A) 3 CG beads at heavy atoms of P, C4’, and N1 for pyrimidine (or P, C4’, and N9 for purine); (B) 2 CG beads at heavy atoms of P and C4’; and (C) 1 CG beads at heavy atom of C4’. Please see also Table 1.

### Residue–separation-based CG statistical potentials

Since RNA folding is generally hierarchical (80), the different residue-separation-ranged interactions may play different roles in stabilizing RNA 3D structures (49). Here, residue separation is described by *k*= |*m*−*n*|, where *m* and *n* stand for the observed nucleotide indices of a pair of atoms along an RNA sequence. In analogy to the newly developed all-atom rsRNASP (77), in cgRNASP, the total energy for an RNA conformation *C* of a given sequence *S* is composed of short-ranged and long-ranged contributions:

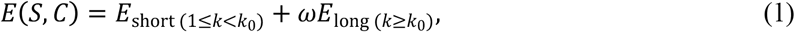

where *k*_0_ is a residue separation threshold to distinguish short- and long-ranged interactions and *ω* is a weight to balance the two contributions.

The long-ranged energy *E*_long_ can be given by (77)

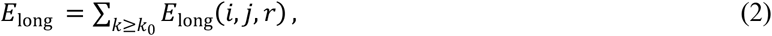

and

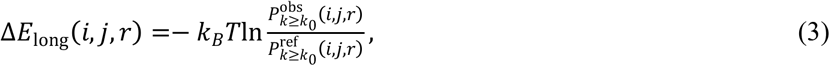

where 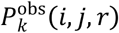 and 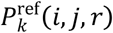 are the probabilities of the distance between atom pairs of types *j* and *k* in distance interval (*r, r* + *dr*] and in the range of residue separation *k*≥*k*_0_ for the native and reference states, respectively.

The short-ranged energy *E*_short_ in cgRNASP is treated in a more complete and subtle way than the all-atom rsRNASP (77) and is given by

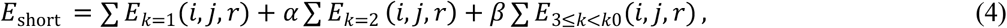

and

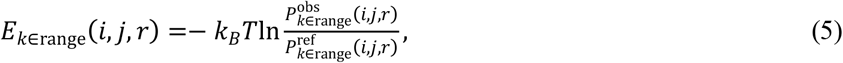

where *k* ∈ range means that residue separation *k* should in the *k* ranges in Eq. (4). 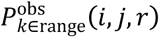 and 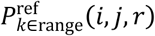 are the probabilities of the distance between atom pairs of types *j* and *k* in distance interval (*r, r* + *dr*] and in the range of residue separation *k*∈range (in Eq. 4) for the native and reference states, respectively. It is noted that beyond the all-atom rsRNASP, the interactions between nearest neighbor residues and between next-nearest neighbor ones were explicitly added in the short-ranged interaction in cgRNASP.

In analogy to the all-atom rsRNASP (77), the average reference state was used for short-ranged energy *E*_short_, while for long-ranged energy *E*_long_, the finite-ideal-gas reference state (59) was used in cgRNASP instead of the random-walk-chain reference state in rsRNASP since the former one also has relatively good performance and has been widely used in protein structure evaluation (48,72). Please see the Supplementary Materials for the details of the use of the reference states for deriving the short- and long-ranged potentials, and see Refs (54,59,72) for the detailed description of the existing reference states.

### Training set and parameters

In cgRNASP, we used the same non-redundant training native set recently used in deriving the all-atom rsRNASP and the dataset is available at https://github.com/Tan-group/rsRNASP(77). The dataset was produced based on the RNA 3D Hub non-redundant set (81) (Release 3.102), which can be found at http://rna.bgsu.edu/rna3dhub/nrlist. The training native set contains 191 RNA structures with chain length >10nt, sequence identity > 80%, coverage > 80% and X-ray resolution < 3.5 Å, and excludes those RNAs complexed with proteins (77). The PDB IDs of these 191 RNAs were also listed in Table S1 in the Supplementary Material. It is noted that a few of RNAs in the test sets have sequence identity > 80% and coverage > 80% with the RNAs in the training native set, and for maintaining the complete structure spectrum, we still kept these RNAs in the training native set (69). For these RNAs in test datasets, the leave-one-out/jackknife method was used in examining the performance of cgRNASP (69, 77).

To optimize the weights (*α, β, ω*) in Eqs. (1) and (4) for short- and long-ranged interactions, we used a training decoy dataset previously built for deriving the all-atom rsRNASP (77), which can be downloaded at https://github.com/Tan-group/rsRNASP. Such training decoy set was generated through four RNA 3D structure prediction models (FARFAR2 (12), RNAComposer (17), SimRNA (43) and 3dRNA v2.0 (21)) with given native secondary structures(82), and contains 35 single-stranded RNAs with wide structure spectrum with about 40 decoy structures for each RNA. In cgRNASP, an RNA length *N*-dependent function *f*(*N*) was also involved to normalize the *N*-dependent CG atom-pair number of the long-ranged interactions due to the large residue-separation range and the consequent *N*-dependent CG atom-pair number (77), and consequently, *ω* in Eq. (1) is equal to *ω*=*ω*_0_/*f*(*N*).

According to the all-atom rsRNASP, *k*_0_ was taken as 5 (77). Based on the examinations on the training decoy dataset, α, *β* and *ω*_0_ were determined for cgRNASP, cgRNASP-PC, and cgRNASP-C, respectively. Please see Section S2 and Fig. S4 in the Supplementary Material for the details of *f*(*N*) and the parameters of α, *β* and *ω*_0_.

In cgRNASP, the distance bin width was taken as 0.3 Å (70, 72, 77), and the distance cutoffs for the potentials in *k* ranges of 1, 2, 3-4, and *k*≥5 were set to the values according to the distance distributions between CG beads in the respective residue separation ranges; see the Supplementary Material for details. For the situation that some atom pairs were not observed within a certain bin width, the potentials were set to the highest potential value in the whole range for corresponding CG atom pair types, and *k*_*B*_*T* was taken as the unit of potential energy.

### Test datasets

To examine the performance of cgRNASP (at different CG levels) and make comparisons with other existing statistical potentials/scoring functions, extensive typical test datasets were used including MD, NM, PM and Puzzles datasets generated by different methods.

Test set_MD consisting of 5 RNAs was generated by Bernauer et al (68) through replica-exchange molecular dynamics simulations with atom position restrained and the RMSDs of the decoy structures are mainly distributed in a narrow range of 0-3 Å (72); Test set_NM composed of 15 RNAs was also generated by Bernauer et al (68) through normal mode perturbation method and the RMSDs of the decoy structures are also mainly distributed in a narrow range of 1-5 Å (72); Test set_PM consisting of 20 RNAs was generated recently by us through four RNA 3D structure prediction models (FARFAR2 (12), RNAComposer (17), SimRNA (43) and 3dRNA v2.0 (21)) with given native secondary structures, and Test set_PM can be downloaded from https://github.com/Tan-group/rsRNASP(77); Test set_Puzzles containing 22 RNAs was obtained from the RNA-Puzzles, which is a CASP-like competition of blind 3D RNA structure predictions, and can be downloaded from https://github.com/RNA-Puzzles/standardized_dataset(74). Test sets PM and Puzzles can be regarded as two realistic test datasets since they are composed of decoy structures of large RNAs generated from different 3D structure prediction models and the RMSDs of decoy structures are distributed in a wide range of ∼2-34 Å (72). The Puzzles dataset is of particular importance since it was generated from the blind CASP-like 3D RNA structure predictions from various research groups with given sequences (74).

### Measuring RNA structure similarity

The metrics of DI (deformation index) was used to measure the structural similarity between two all-atom RNA structures, and DI is defined as (83):

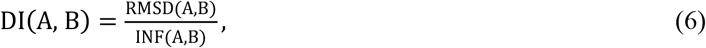

where RMSD(A, B) and INF(A, B) reflects the difference of geometry and topology between structures A and B, respectively. RMSD(A, B) is the root-mean-square-deviation (RMSD) between structures A and B. INF(A, B) is the interaction network fidelity between structures A and B, and is measured by Matthews correlation coefficient of base-pairing and base-stacking interactions (84). The tools for calculating DI and INF can be downloaded from https://github.com/RNA-Puzzles/BasicAssessMetrics(85).

## Results and discussion

In the following, we examined the performance of cgRNASP against extensive test datasets. As described in the subsection of test datasets, test sets MD and NM were mainly generated from perturbation methods, while test sets PM and Puzzles generated from various 3D structure prediction models can be considered as realistic test datasets and the Puzzles dataset from the blind CASP-like RNA structure predictions is of particular importance. Therefore, we first examined the overall performance of 3-bead cgRNASP against all the test sets, and afterwards focused on the performance of cgRNASP at different CG levels and the computation efficiency of cgRNASP against the realistic RNA-Puzzles dataset.

## Evaluation metrics

To describe the performance of cgRNASP, in analogy to previous works (71,72,77), we used the number of native structures with the lowest energy, the DI of lowest-energy structure (including native one), and the DI of lowest-energy decoy structure (excluding native one) as evaluation metrics for identifying native/near-native structures. Moreover, we used Pearson correlation coefficient (PCC) (69,77) as an evaluation metrics for ranking decoy structures, and PCC for decoy structures of an RNA can be calculated as (69):

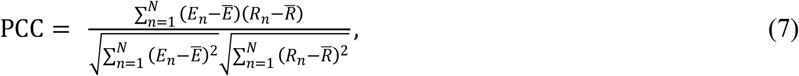

where *E*_*n*_ and *R*_*n*_ are the energy and DI of the *n*th decoy for the RNA, respectively. Ē-and 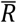 -are the average energy and DI of all decoys for the RNA, respectively. The value of PCC ranges from 0 to 1, and PCC of 1 represents a perfect performance of the statistical potential.

## Overall performance of cgRNASP on test datasets

### Overall performance of cgRNASP on all the test datasets

As shown in Fig.2A-G and Table S2 in the Supplementary Material, for all the test sets including the test sets MD and NM with small RNAs from perturbation methods and the realistic test sets PM and Puzzles with large RNAs from various 3D structure prediction models, cgRNASP identifies ∼77% native structures, i.e., 48 native structures out of decoy ones of 62 RNAs. Such values are identical to those of the newly developed all-atom rsRNASP (∼77% and 48 out of 62 RNAs), and appears higher than the all-atom statistical potentials/scoring functions of RNA3DCNN (∼74% and 46 out of 62), ARES (∼11% and 7 out of 62), DFIRE-RNA (∼56% and 35 out of 62), 3dRNAscore (∼34% and 21 out of 62), and RASP (∼26% and 16 out of 62). This indicates that cgRNASP identifies similar number of native structures with rsRNASP while more native ones than other all-atom statistical potentials/scoring functions for all the test datasets. Furthermore, we calculated the mean DI of lowest-energy structures including native ones and that excluding native ones. As shown in Figs. 2B and C, the mean DI of lowest-energy structures and that excluding native structures calculated from cgRNASP are 1.8 Å and 7.0 Å, which are almost identical to those from the all-atom rsRNASP (2.1 Å and 6.9 Å). However, such two values from cgRNASP are both visibly smaller than those from other all-atom statistical potential/scoring functions of RNA3DCNN (3.1 Å and 8.3 Å), ARES (9.3 Å and 9.5 Å), DFIRE-RNA (4.8 Å and 7.8 Å), 3dRNAscore (8.3 Å and 9.6 Å), and RASP (9.1 Å and 10.4 Å), suggesting that cgRNASP identifies the structures more similar to native ones.

**Figure 2.**
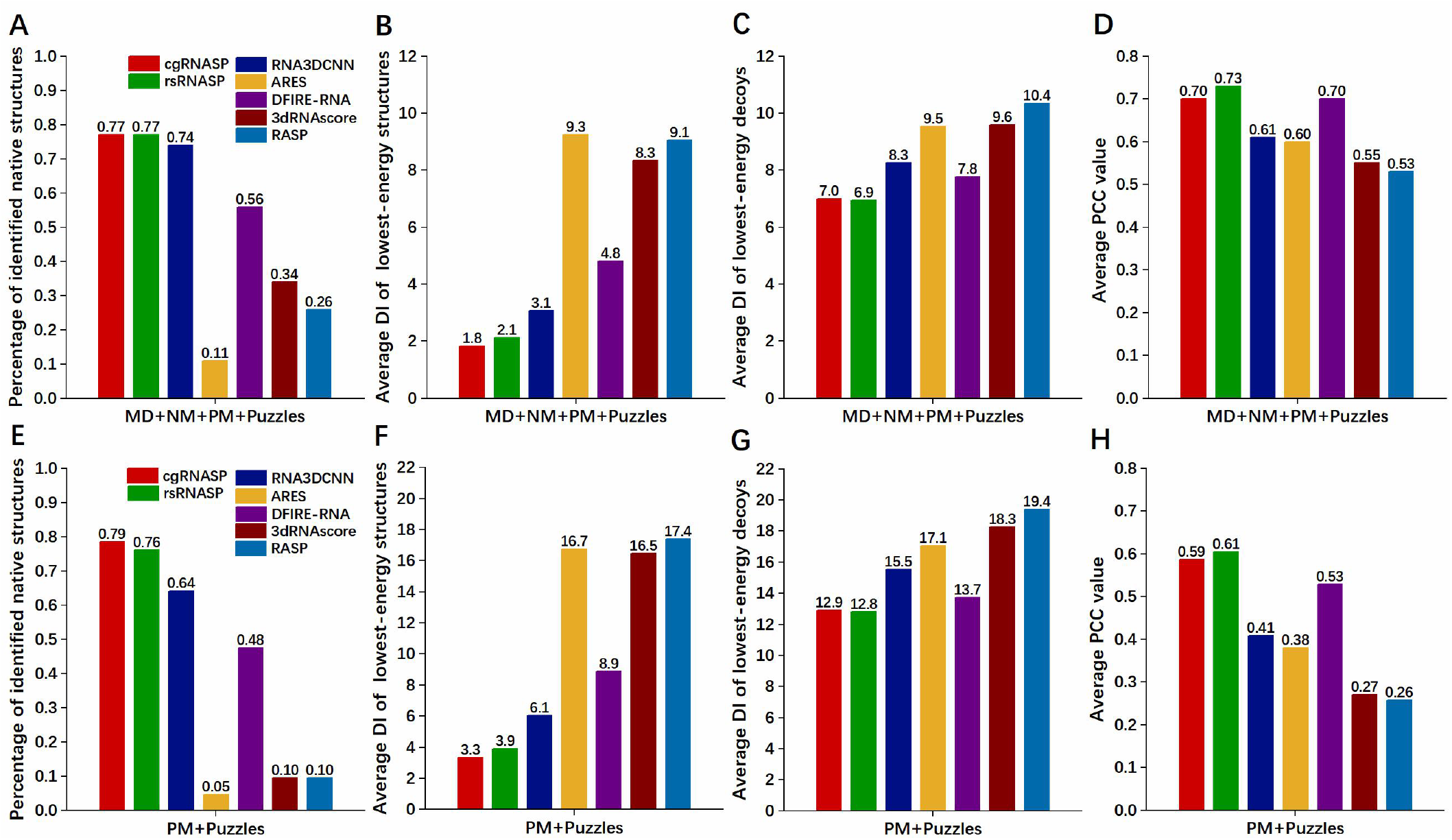
(A,E) Percentages of the number of identified native structures, (B,F) average DI values of structures with the lowest energy (including native ones), (C,G) average DI values of decoy structures with the lowest energy (excluding native ones), and (D,H) average PCC values between DIs and energies by coarse-grained cgRNASP and other all-atom statistical potentials. Panels (A-D) are for all the test sets (MD+NM+PM+Puzzles), and (E-H) are for the realistic test sets (test sets PM+Puzzles) from various 3D structure prediction models. Here, the PCC values for the whole test sets (MD+NM+PM+Puzzles) and for the realistic sets (PM+Puzzles) were averaged over the mean values of respective test sets since decoys in a test set were generated with the same method and thus have similar structure features.

To examine the performance of cgRNASP in ranking decoy structures, we calculated the PCC values between energies and DIs for all the test datasets by cgRNASP and other all-atom statistical potentials/scoring functions; see Eq. (7). As shown in Fig. 2D for all the test sets, cgRNASP has an overall comparable performance to the all-atom rsRNASP in ranking decoy structures since the average PCC value (0.7) from cgRNASP is very slightly smaller than that (0.73) from all-atom rsRNASP. Furthermore, the PCC value from cgRNASP is identical to that (PCC ∼0.7) from the all-atom DFIRE-RNA while is apparently larger than other all-atom statistical potential/scoring function of RNA3DCNN (PCC∼0.61), ARES (PCC∼0.60), 3dRNAscore (PCC∼0.55) and RASP (PCC∼0.53), respectively. Namely, for all the test datasets, cgRNASP has similar performance to the all-atom DFIRE-RNA while apparently better performance than other all-atom statistical potentials in ranking decoy structures.

Therefore, for all the test datasets, the present (coarse-grained) cgRNASP has an overall similar performance with the newly developed all-atom rsRNASP while is overall superior to other existing all-atom statistical potential/scoring functions in identifying RNA native/near-native structures and in ranking decoy structures.

### Performance of cgRNASP on realistic test datasets-PM and puzzles

Test datasets PM and Puzzles generated from various RNA 3D structure prediction models contains large RNA decoy structures with large RMSD range, and consequently can serve as realistic test sets for a statistical potential beyond test sets MD and NM composed of small RNAs and near-native decoy structures from perturbation methods; see subsection of test datasets.

As shown in Figs. 2E-G and Table S2 in the Supplementary Material, for the PM and Puzzles datasets, cgRNASP identified 79% native structures (33 out of 44), and the mean DI of the lowest-energy structures and that excluding native ones from cgRNASP are 3.3 Å and 12.9 Å, respectively. Such three values are overall very slightly superior to those from the all-atom rsRNASP (∼76%, 3.9 Å, and 12.8 Å), suggesting a very slightly better performance of cgRNASP than the all-atom rsRNASP in identifying native/near-native structures for test datasets PM and Puzzles. However, the comparison of the three metrics values with other all-atom statistical potentials/scoring functions indicates that cgRNASP has apparently better performance than RNA3DCNN (∼64%, 6.1 Å, and 15.5 Å), ARES (∼5%, 16.7 Å, and 17.1 Å), DFIRE-RNA (∼48%, 8.9 Å, and 13.7 Å), 3dRNAscore (∼10%, 16.5 Å, and 18.3 Å) and RASP (∼10%, 17.4 Å, and 19.4 Å).

Moreover, as shown in Fig. 2H for the PM and Puzzles datasets, the PCC value (0.59) from cgRNASP is very close to that (0.61) from rsRNASP while is apparently higher than other all-atom statistical potentials/scoring functions of RNA3DCNN (0.41), ARES (0.38), DFIRE-RNA (0.53), 3dRNAscore (0.27) and RASP (0.26). This suggests that cgRNASP has the same performance with the all-atom rsRNASP and apparently better performance than other all-atom statistical potentials/scoring functions in ranking decoy structure for the PM and Puzzles datasets.

Therefore, overall, for realistic datasets including PM and Puzzles, the present coarse-grained cgRNASP has very slightly better performance than the newly developed all-atom rsRNASP in identifying native/near-native structures, while appears apparently superior to other all-atom statistical potentials/scoring functions in identifying native/near-native structures and in ranking decoy ones.

## Performance of cgRNASP on Puzzles dataset

Test dataset Puzzles composed of 22 RNAs has been widely considered as a realistic test dataset and consequently is of particular importance since it was generated from the blind CASP-like 3D RNA structure predictions from various top research groups with given sequences (74).

As shown in Figs. 3A-C and Table S3 in the Supplementary Material, for the Puzzles dataset, cgRNASP identifies 18 native structures out from decoys of 22 RNAs, and the mean DI of lowest-energy structures and that excluding native ones from cgRNASP are 1.5 Å and 13.7 Å. In contrast, rsRNASP identifies 16 native ones out from decoys of 22 RNAs and the mean DI of lowest-energy structures and that excluding native ones from rsRNASP are 4.6 Å and 14.4 Å, respectively. Moreover, the three metrics values from cgRNASP appear apparently better than those from other statistical potential/scoring functions of RNA3DCNN (13, 5.9 Å, and 18.5 Å), ARES (2, 18.1 Å, and 18.8 Å), DFIRE-RNA (10, 7.6 Å, and 14.4 Å), 3dRNAscore (2, 17.1 Å, and 19.4 Å) and RASP (2, 17.8 Å, and 20.0 Å). Thus, for the Puzzles dataset, cgRNASP is slightly superior to rsRNASP and has apparently better performance than other all-atom statistical potentials/scoring functions in identifying native/near native structures.

**Figure 3.**
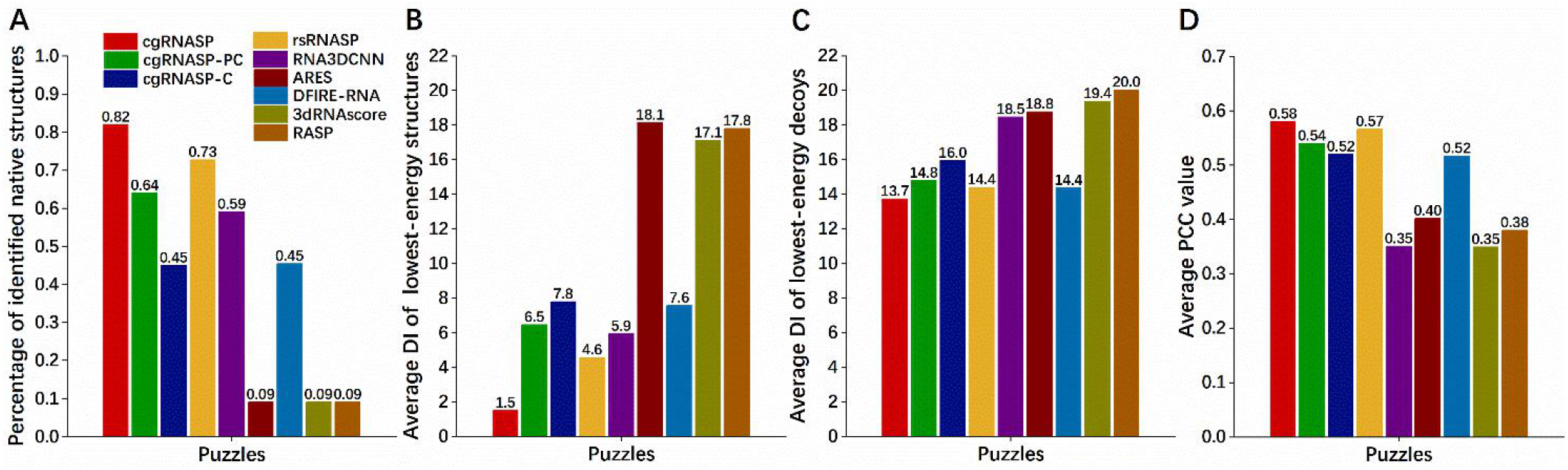
(A) Percentages of the number of identified native structures, (B) average DI values of structures with the lowest energy (including native ones), (C) average DI values of decoy structures with the lowest energy (excluding native ones), and (D) average values of PCCs between DIs and energies by cgRNASP at different CG levels (cgRNASP, cgRNASP-PC, and cgRNASP-C) and other all-atom statistical potentials for the test set Puzzles.

Furthermore, as shown in Fig. 3D for the Puzzles dataset, the PCC between DIs and energies calculated from cgRNASP is 0.58, which is slightly higher than that from rsRNASP (0.57). Moreover, such value of cgRNASP is apparently higher than those from other all-atom statistical potentials/scoring functions of RNA3DCNN (0.35), ARES (0.40), DFIRE-RNA (0.52), 3dRNAscore (0.35) and RASP (0.38). Thus, for the Puzzles dataset, cgRNASP is slightly better than rsRNASP and is apparently superior to other all-atom statistical potentials/scoring functions in ranking decoy structures.

Therefore, for the Puzzles dataset from CASP-like blind 3D structure prediction competition, our present coarse-grained cgRNASP is slightly better than the all-atom rsRNASP and is apparently superior to the other all-atom statistical potentials/scoring functions in identifying native/near-native structures and in ranking decoy ones.

## Performance of cgRNASP at different CG levels on Puzzles dataset

In this subsection, we would examine the relative performance of cgRNASP at different CG levels on the Puzzles dataset, i.e., 3-bead cgRNASP, 2-bead cgRNASP-PC, and 1-bead cgRNASP-C; see Materials and Methods.

As shown in Figs. 3A-C, cgRNASP, cgRNASP-PC, and cgRNASP-C identify 18, 14, and 10 native structures out of decoy ones of 22 RNAs in the Puzzles dataset, respectively. The mean DI with the lowest-energy structures and that excluding native ones are 1.5 Å and 13.7 Å for cgRNASP, 6.5 Å and 14.8 Å for cgRNASP-PC, and 7.8 Å and 16.0 Å for cgRNASP-C, respectively. Thus, in identifying native/near-native structures, the performance of cgRNASP at different CG levels follows the order of cgRNASP>cgRNASP-PC>cgRNASP-C for the Puzzles dataset, which is consistent with the number of CG beads used in the respective potentials. Furthermore, Fig. 3D shows that the PCC values between energies and DIs of decoys are 0.58, 0.54, and 0.52 for cgRNASP, cgRNASP-PC, and cgRNASP-C, respectively. Thus, the performance in ranking decoy structures in the Puzzles dataset follows the order of cgRNASP>cgRNASP-PC> cgRNASP-C, which is identical to the above order in identifying native/near-native structures. It is understandable that the performance of cgRNASP follows the order of cgRNASP>cgRNASP-PC> cgRNASP-C, since more CG beads would generally involve more geometric information and constraints for a structure and consequently would be more effective for structure evaluation.

Overall, compared with other all-atom statistical potentials/scoring functions, for the realistic Puzzles dataset, cgRNASP has slightly better performance with mean DI of lowest-energy structures ∼1.5 Å and PCC∼0.58 than the all-atom rsRNASP (with the two values of 4.6 Å and 0.57). Moreover, cgRNASP-PC has also acceptable performance with mean DI of lowest-energy structures ∼6.5 Å and PCC∼0.54 overall beyond existing all-atom statistical potentials/scoring functions except for rsRNASP. Additionally, it is encouraging that cgRNASP-C based on extremely reduced 1-bead representation of C4’ atom has very similar performance to the all-atom DIRE-RNA and appears superior to the all-atom RNA3DCNN in ranking decoys.

## Evaluation efficiency of cgRNASP on Puzzles dataset

A reliable CG-based statistical potential can be used not only for evaluating CG structures but also for evaluating all-atom structures at high efficiency due to the greatly reduced atom representation. Here, we would examine the computation efficiency of cgRNASP on the Puzzles dataset, in a comparison with existing top all-atom statistical potentials/scoring functions.

As shown in Fig. 4, for evaluating decoys of RNAs in the Puzzles dataset, cgRNASP is strikingly more efficient than the all-atom statistical potentials/scoring functions of rsRNASP, RNA3DCNN and DFIRE-RNA. Specifically, for the Puzzles dataset, the mean computation time of (3-bead) cgRNASP is only ∼1/65 of that of rsRNASP, while rsRNASP has comparable computation time with DFIRE-RNA and is ∼10 times more efficient than RNA3DCNN. Furthermore, the computation time of cgRNASP slightly decrease with the decrease of number of CG beads used in cgRNASP. For example, cgRNASP is ∼2 times less efficient than cgRNASP-PC and is ∼9 times less efficient than cgRNASP-C. It is understandable that cgRNASP is strikingly more efficient than all-atom statistical potentials/scoring functions and cgRNASP with less CG beads is more efficient since the computation time of a statistical potential should be approximately proportional to the square of the number of (CG) atoms per nucleotide involved in the potential.

**Figure 4.**
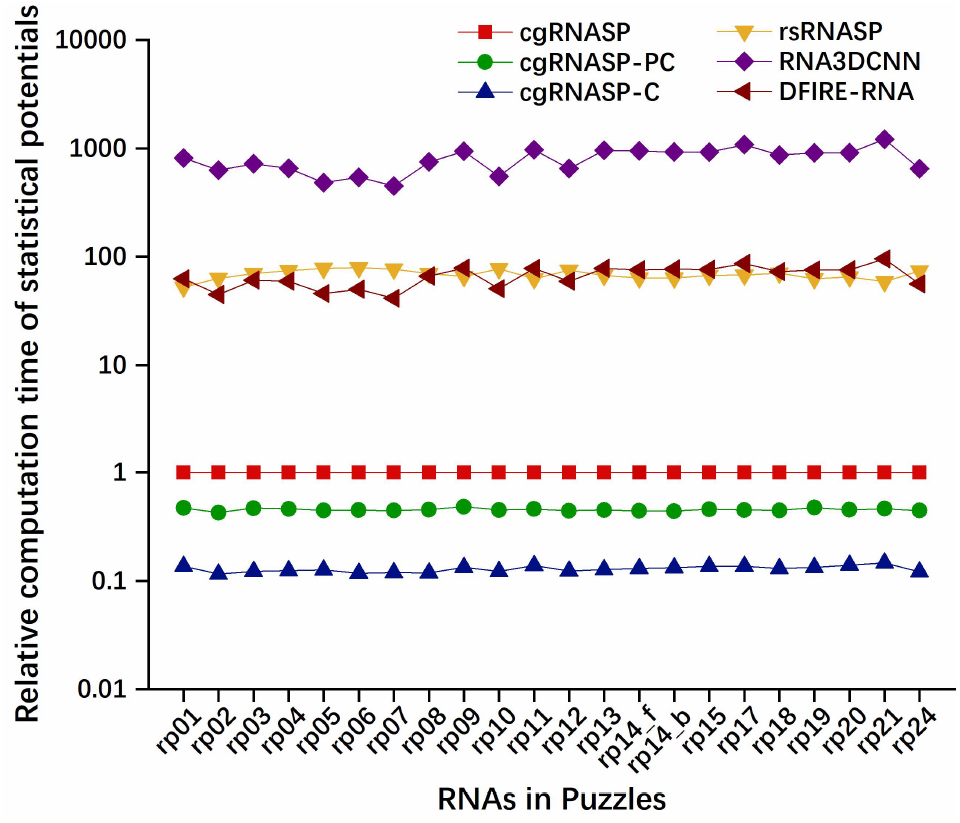
Computation times of cgRNASP at different CG levels (cgRNASP, cgRNASP-PC, and cgRNASP-P) and other top all-atom statistical potentials for the RNAs in the test set Puzzles, relative to that of 3-bead cgRNASP. Here, rp14-f and rp14-b stand for rp14-free and rp14-bound, respectively.

The strikingly higher efficiency of cgRNASP with good performance would be very beneficial to RNA 3D structure evaluation through either greatly saving evaluation time or evaluating much more structure candidates within a given time.

## How can cgRNASP with coarse-grained representation have good performance?

As shown above, the present (3-bead) cgRNASP with coarse-grained representation can have similar performance for all the test dataset and even slightly better performance for the Puzzles dataset, compared with the newly developed all-atom rsRNASP. Then how can cgRNASP have good performance with highly reduced (coarse-grained) representation?

First, we examined the individual contributions of short- and long-ranged interactions against the Puzzles dataset. As shown in Figs. 5A-D, in identifying native/near-native structures and in ranking decoy ones, the long-ranged contribution in cgRNASP performs comparable to that in rsRNASP while the short-ranged contribution with 3≤*k*≤4 in cgRNASP performs worse than that in rsRNASP; see also Fig. 3 in Ref (77). However, the explicit involvement of the potentials with *k*=1 and *k*=2 in cgRNASP causes an overall comparable short-ranged contribution to that in rsRNASP (77). The combination of long-ranged and short-ranged potentials with comparable performance to those of rsRNASP leads to the overall similar performance of cgRNASP to that of rsRNASP. It is not very strange that cgRNASP is very slightly superior to rsRNASP for the Puzzles dataset, which should be attributed to the more complete/subtle involvement of short-ranged potentials at different residue separation since the all-atom rsRNASP did not involve the interactions at residue separations with *k*=1 and *k*=2. The overall similar performance of cgRNASP to rsRNASP may also be attributed to the 3-bead CG representation in cgRNASP which can be considered a minimal one including both backbone and base.

**Figure 5.**
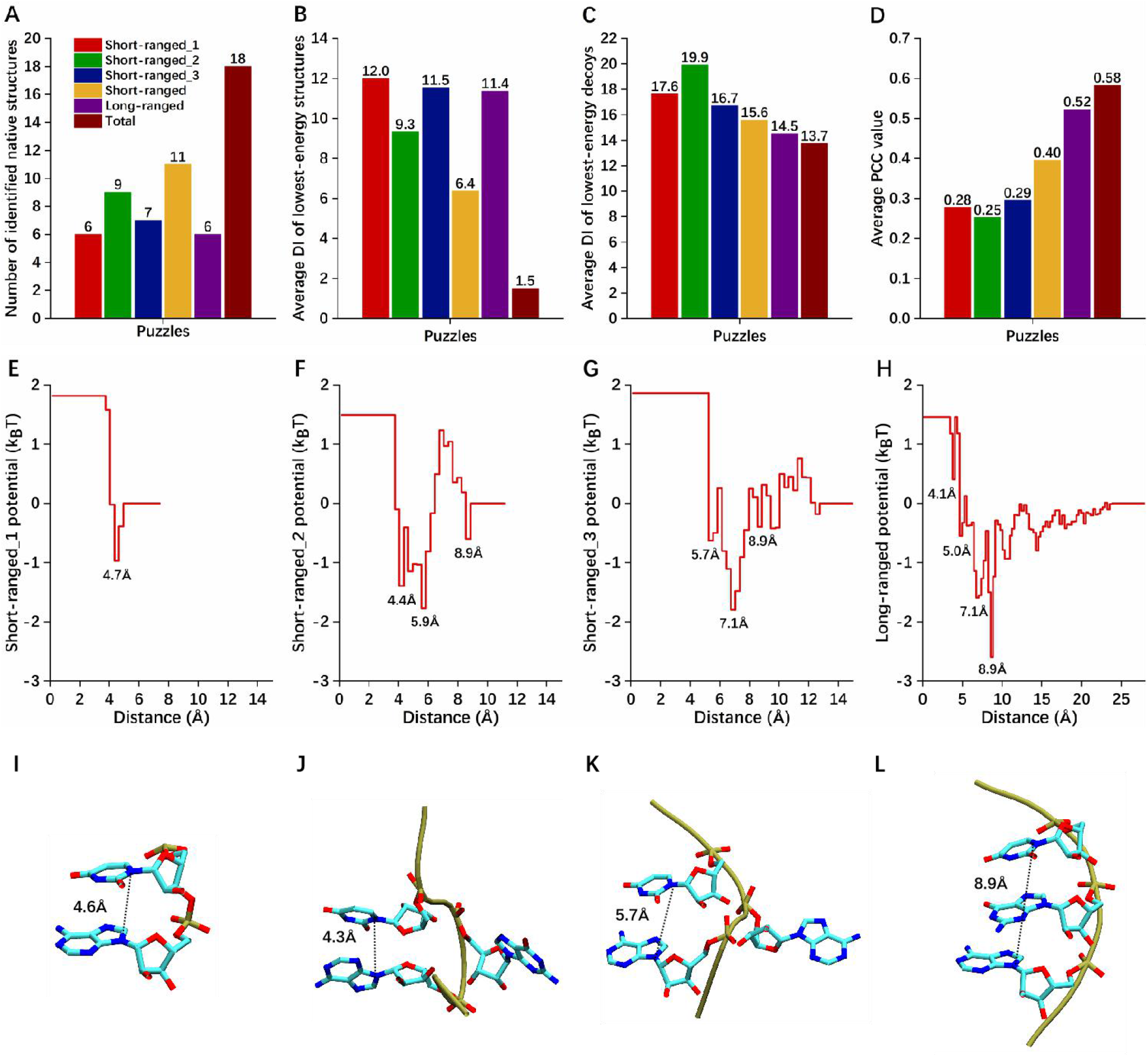
(A) Percentages of the number of identified native structures, (B) average DI values of structures with the lowest energy (including native ones), (C) average DI values of decoy structures with the lowest energy (excluding native ones), and (D) average values of PCCs on DIs by short-ranged and long-ranged energies from cgRNASP. (E-G) Short-ranged and (H) long-ranged potentials between AN9 and UN1 in cgRNASP. In panels (A-H), short-ranged_1, short-ranged_2, and short-ranged_3 potentials denote those at residue separations of *k*=1, *k*=2, and 3≤*k*≤4, respectively. (I-J) Representative distances between AN9 and UN1 for nearest-neighbor base stacking (I), next-nearest neighbor U-turns (J and K), and next-nearest neighbor base-stacking (L) captured in the short-ranged potentials. Here, we did not show other representative distances for reverse Hoogsteen (∼7.1 Å) and Watson-Crick base pairing (∼8.9 Å) in the short-ranged_3 potential, as well as those for base stacking between adjacent branches (∼4 Å), base stacking at triplex (∼5 Å) and coaxial-stacking at junctions (∼5 Å) in the long-ranged potential, respectively, which have been illustrated in the Supplementary Material and Ref [73].

Furthermore, we examined the microscopic interactions captured in cgRNASP, in a comparative way with rsRNASP (77). As shown in Figs. 4E-F for the potentials between N9 atom of adenine (A) and N1 atom of uracil (U) as a paradigm, the long-ranged potential in cgRNASP captures similar minimums with that in rsRNASP while the short-ranged potentials can capture more minimums than that in rsRNASP; see also Figs. 4AB in Ref (77). This is consistent with the above discussed performances of short- and long-ranged contributions in cgRNASP. Compared with rsRNASP (77), the explicit and complete involvement of short-ranged potentials at different residue separations leads to more detailed inter-bead (atom) interactions captured in the short-ranged potential including nearest-neighbor base stacking (at ∼4.6 Å), next-nearest-neighbor U-turns (at ∼4.3 Å and ∼5.7 Å), and next-nearest-neighbor base stacking (at ∼8.9 Å), in addition to the interactions captured in rsRNASP such as canonical Watson-Crick base pairing (at ∼8.9 Å), the intra-loop base interactions (at ∼5.7 Å), and the reverse Hoogsteen base-pairing interaction (at ∼7.1 Å) (77). Moreover, similar to rsRNASP (77), the long-ranged potential in cgRNASP also captures some tertiary base-stacking interactions including base-stacking between two residues in two adjacent branches of RNA 3D structures (e.g., tetraloop receptor), coaxial-stacking at junction region and base-stacking in triple-helical region; see Fig. S5 in the Supplementary Material and see also Ref (77).

Therefore, the combination of more detailed short-ranged potential and long-ranged one contribute to the good performance of cgRNASP even with highly reduced (coarse-grained) representation.

## Conclusion

In this work, we have developed a series of residue-separation based CG statistical potentials at different CG levels (cgRNASP) for RNA 3D structure evaluation and cgRNASP can be rather applicable for the CG structure prediction models and has much higher computation efficiency for all-atom structure evaluation than those all-atom statistical potentials and scoring functions. The CG potentials of cgRNASP were built through distinguishing short- and long-ranged interactions, and compared with the all-atom rsRNASP, more complete and subtle treatment on the short-ranged interactions through adding interactions between nearest neighbor residues and between next-nearest neighbor ones. Extensive tests against the available datasets indicate that, overall, cgRNASP has varying performance with different CG levels and can have similarly good or even slightly better performance for the realistic Puzzles dataset, compared with the all-atom rsRNASP. Furthermore, cgRNASP has strikingly higher computation efficiency than all-atom statistical potentials/scoring functions and has overall better performance than other existing traditional statistical potentials and scoring functions from neural networks, especially for the realistic test dataset including the Puzzles dataset. It should be emphasized that our cgRNASP can be directly used for evaluating CG structure candidates generated from existing related CG-based structure prediction models and can be also very useful for evaluating all-atom structure candidates at strikingly higher efficiency than all-atom statistical potentials.

The present CG statistical potentials of cgRNASP can be furtherly improved for more applicable versions to different CG structure prediction models and for more accurate evaluation on RNA 3D structures. First, cgRNASP potentials at different CG levels only involve those real heavy atoms rather than dummy atoms (e.g., mass centers of atom groups). For more specific application, the present cgRNASP can be extended to include those dummy atoms used in some specific CG structure prediction models and such specifically improved potentials can be more useful for those corresponding CG structure prediction models. Second, some other geometric parameters such as torsional angle and orientation can be involved in cgRNASP to enhance the description of the relative relationship between atoms or atom groups beyond pair atom-atom distance (48,70,86). Third, a multi-body (three-or four-body) potential can be explicitly supplemented to cgRNASP which would improve the description on correlated atom-atom distributions (48,87). Fourth, the reference states in cgRNASP can be possibly circumvented through some treatments such as iterative technique since the used reference states may still deviate from the ideal one (65,66). Fifth, due to the currently limited RNA native structures in PDB database (7), cgRNASP can be continuously improved with the increase of the number of experimentally determined RNA native structures. Finally, some underlying geometric information may be captured from training through neural network over high-quality training dataset, and the captured information may be combined to cgRNASP to derive a new CG statistical potential/scoring function with higher performance.

Nevertheless, the present CG statistical potentials of cgRNASP would be definitely very beneficial to related CG-based 3D structure evaluations, as well as to all-atom-based 3D structure evaluations at strikingly high computation efficiency.

## Data availability Statement

All relevant data are within the paper and its Supporting Information files. The potentials of cgRNASP at different CG levels are available at website https://github.com/Tan-group/cgRNASP.

## Supporting information

Supplemental Material is available for this article.

## Acknowledgments

We are grateful to Profs Shi-Jie Chen (University of Missouri) and Jian Zhang (Nanjing University) for valuable discussions. The numerical calculations in this work were performed on the super computing system in the Super Computing Center of Wuhan University.

## Author Contributions

ZJT, BZ and YLT designed the research. YLT, XW and SXY performed the research. ZJT, YLT, XW and BZ analyzed the data. YLT, XW, BZ and ZJT wrote the manuscript.

## Funding

This work was supported by grants from the National Science Foundation of China (12075171, 11774272).

## Notes

https://github.com/Tan-group/cgRNASP

### Competing Interest Statement

The authors have declared no competing interest.

