## Supplementary Material for "cgRNASP: coarse-grained statistical potentials with residue separation for RNA structure evaluation"

### 1. A detailed description of the statistical potential of cgRNASP

Our present cgRNASP potentials were developed based different coarse-grained (CG) levels, which has been described in details in the main text. Similar to the recently developed rsRNASP (1), cgRNASP is composed of short- and long-ranged energy functions distinguished by residue separation *k*, which is defined by $k = |m - n|$, where *m* and *n* correspond to the observed residue sequence positions of a pair of atoms along an RNA chain. Compared with the newly developed all-atom rsRNASP (1), the short-ranged interaction in cgRNASP was involved more subtly and completely through explicitly adding the interactions between nearest neighbor residues and between next-nearest ones.

For extracting short-ranged interactions at the reside separations of *k*=1, *k*=2, and $3 \leq k < k_0$, the averaging reference state was used (2). Besides, to avoid the problem of sparse data in short-ranged potential, we employed the method developed by Sippl (3). Thus, the short-ranged potential can be given by

$$\Delta E_{\text{short}, k\in\text{range}}(i,j,r) = k_B T \ln\left[1 + M_{ij}\sigma\right] - k_B T \ln\left[1 + M_{ij}\sigma \frac{P^{\text{obs}}_{k\in\text{range}}(i,j,r)}{P^{\text{obs}}_{k\in\text{range}}(r)}\right], \quad (1)$$

where $k \in \text{range}$ stand for the residue separations of *k*=1, *k*=2, and $3 \leq k < k_0$ in the short range and $k_0$ is the residue separation threshold to distinguish short- and long-ranged interactions. $P^{\text{obs}}_{k\in\text{range}}(i,j,r)$ is the observed probability of the distance between atom pair of atom types $i$ and $j$ in distance interval $(r, r + dr]$ within residue separation interval $k \in \text{range}$, and $P^{\text{obs}}_{k\in\text{range}}(r)$ is the summation of $P^{\text{obs}}_{k\in\text{range}}(i,j,r)$ over all kinds of atom pair types. $M_{ij}$ is the total number atom pair of atom types *i* and *j* observed within residue separation intervals $k \in \text{range}$. $\sigma$ was set to 0.02, as proposed by Sippl (3).

For long-ranged potential, the finite-ideal-gas reference state was used (4). Thus, according to previous description (5), the probability of reference state $P^{\text{ref}}_{k\geq k_0}(i,j,r)$ can be given by (5):

$$P^{\text{ref}}_{k\geq k_0}(i,j,r) = \frac{N^{\text{obs}}_{k\geq k_0}(i,j,r_c)}{N^{\text{obs}}_{k\geq k_0}} \left(\frac{r}{r_c}\right)^D = P^{\text{obs}}_{k\geq k_0}(r_c)\left(\frac{r}{r_c}\right)^D, \quad (2)$$

where $P^{\text{obs}}_{k\geq k_0}(i,j,r_c)$ is the observed probability of the distance between atom pair of atom types *i* and *j* within residue separation interval $k \geq k_0$ at distance cutoff $r_c$ for training native set. Here, a dimension parameter $D$ is involved since macromolecule systems are actually not ideal gas even at high temperature, and according to Ref (6), *D* was taken as 2.0 for cgRNASP, cgRNASP-PC and cgRNASP-C. Thus, the long-ranged potential can be given by:

$$\Delta E_{\text{long}}(i,j,r) = - k_B T \ln \frac{P^{\text{obs}}_{k\geq k_0}(i,j,r)}{P^{\text{obs}}_{k\geq k_0}(r_c)\left(\frac{r}{r_c}\right)^D}. \quad (3)$$

Therefore, in cgRNASP, the total energy for an RNA conformation $C$ of a given sequence $S$ is composed of short-ranged and long-ranged contributions:

$$E(S,C) = E_{\text{short}\,(1<k<k_0)} + \omega E_{\text{long}\,(k_0\leq k)}, \quad (4)$$

where $E_{\text{short}\,(1<k<k_0)}$ is given by

$$E_{\text{short}} = \sum E_{k=1}(i,j,r) + \alpha \sum E_{k=2}\,(i,j,r) + \beta \sum E_{3\leq k<k0}(i,j,r). \quad (5)$$

**2. Determination of the parameters involved in cgRNASP at different CG levels**

The residue separation threshold $k_0$ was introduced to distinguish short- and long-ranged interactions, and $k_0$ describes the residue separation boundary between intra-loop interactions and those beyond in RNA structures. According to rsRNASP (1), the residue separation threshold $k_0$ was taken as 5 to distinguish short- and long-ranged interactions in cgRNASP. According to the distribution of distance between bead pairs for different residue separations ($k \geqslant k_0$, $k_0 > k \geqslant 3$, $k$=2 and $k$=1) from training native set and Ref (1), the distance cutoff $r_c$ for the long-ranged potential, the short-ranged potential at $k_0 > k \geqslant 3$, the short-ranged potential at $k$=2, and that at $k$=1 were taken as 24 Å, 13 Å, 9 Å, and 5 Å for cgRNASP and cgRNASP-PC, and 24 Å, 16 Å, 12 Å, and 7 Å for cgRNASP-C; see Fig. S1. Here, the values of $r_c$ of cgRNASP-C are slightly larger than those of cgRNASP and cgRNASP-PC since the distance separation between beads of C4' atoms in cgRNASP-C is generally larger than those between more CG beads of cgRNASP and cgRNASP-PC in RNA structures.

Different from short-ranged interactions limited by much less residue separation and shorter distance cutoff, for a CG bead, the number of its bead pairs of long-ranged interactions could strongly depend on RNA length $N$ (in nt). Thus, at first, an $N$-dependent function is required to normalize the energy contribution from the long-ranged interaction for cgRNASP, similar to rsRNASP (1). The relations between the numbers of long-ranged bead pairs within 24 Å scaled by $N$ and RNA length $N$ from the statistical analyses on the training native set has been shown in Fig. S3 for different CG levels. According to Fig. S3, for convenience and simplicity, we employed the following fitted functions in our cgRNASP at different CG levels: $f_1(N) = \frac{-355}{(N+16)^{0.5}} + 72$ for cgRNASP; $f_2(N) = \frac{-290}{(N+29)^{0.5}} + 50$ for cgRNASP-PC; $f_3(N) = \frac{-175}{(N+39)^{0.5}} + 27$ for cgRNASP-C. Therefore, the weight $w$ in cgRNASP can be furtherly given by:

$$w = w_0/f(N). \quad (6)$$

In our cgRNASP, $w_0$, α, and β in Eq. 4 (in the main text) were determined by the training decoy set which was built from the RNA 3D structure prediction models (1) and includes 35 single-stranded RNAs with about 40 decoy structures for each RNA. Specifically, we used two metrics, including the number percentage of native structures identified within top five of the lowest energies and Pearson correlation coefficient (PCC) values on DIs (7) to optimize the parameters of ($w_0$, α, and β) (1). As illustrated in Fig. S5, the proper choice of the values of ($w_0$, α, β) can achieve the compromise performance on these three metrics (number of native structures identified within top five (lowest-energy) structures, average DI of lowest-energy decoy structures, and average PCC value on DIs) for the training decoy set (1, 8-11). Thus, ($w_0$, α, β) were taken as (3.5, 1.0, 1.7) for cgRNASP, (6.2, 0.5, 1.2) for cgRNASP-PC, and (40.0, 3.0, 3.0) for cgRNASP-C, respectively.

Table S1. PDB IDs of 191 RNAs in our training native set for cgRNASP [a].

| | | | | | | | | | | | | |
|---|---|---|---|---|---|---|---|---|---|---|---|---|
| 1CSL | 1D4R | 1DUH | 1DUQ | 1ET4 | 1EVV | 1F1T | 1F27 | 1FIR | 1FUF | 1I9X | 1KD5 | 1KFO |
| 1KH6 | 1KXK | 1L2X | 1MHK | 1NBS | 1NTA | 1NUV | 1Q96 | 1QBP | 1QCU | 1RNA | 1SDR | 1T0E |
| 1U9S | 1XJR | 1Y26 | 1Y27 | 1YFG | 1YLS | 1Z7F | 1ZCI | 205D | 255D | 280D | 283D | 2A64 |
| 2ET8 | 2G92 | 2GDI | 2H1M | 2IL9 | 2JLT | 2O3Y | 2OE8 | 2OEU | 2OIU | 2P7E | 2Q1R | 2QUS |
| 2QWY | 2R1S | 2YGH | 2ZY6 | 353D | 359D | 361D | 364D | 387D | 397D | 3BNN | 3CGS | 3CJZ |
| 3CZW | **3D2V** | 3DIL | 3E5C | 3GM7 | 3GS5 | 3IBK | 3IGI | 3K1V | 3LOA | 3MEI | 3MJA | 3ND3 |
| 3NPQ | 3OXE | 3P22 | 3P59 | 3PDR | **3Q3Z** | **3R4F** | **3RG5** | 3SJ2 | **3SKL** | **3SUX** | 406D | 413D |
| 422D | 433D | 4E48 | 4E5C | **4ENC** | **4FRN** | 4GXY | 4J50 | **4JF2** | **4JRC** | 4JRD | 4JRT | **4K27** |
| **4KQY** | 4KYY | **4LVW** | 4MCF | 4NFQ | 4NLF | 4O41 | 4OQU | 4P3T | 4P95 | 4P97 | 4PCJ | 4PHY |
| **4PLX** | **4PQV** | 4QJD | 4QK9 | 4QLM | 4R4V | 4RBQ | 4RBY | 4RGE | 4RUM | 4RZD | 4TS2 | **4WFL** |
| 4XWF | **4Y1M** | **4YAZ** | **4ZNP** | 5AY2 | 5BJO | **5BTM** | 5BTP | 5C5W | 5CNR | 5EW4 | 5G4T | 5KPY |
| 5KTJ | 5KVJ | 5L4O | **5LYS** | **5M0H** | **5ML7** | 5MWI | 5NDI | 5NWQ | 5NXT | **5OB3** | 5T83 | **5U3G** |
| 5UNE | 5UZ6 | 5V1K | 5V3F | 5VJ9 | 5XWG | 5Z1I | 6C63 | 6CB3 | 6CK5 | 6CU1 | 6D3P | **6DLR** |
| **6DME** | 6DN2 | **6DVK** | 6E1S | 6E7L | **6E8U** | 6FZ0 | **6H0R** | 6HC5 | 6HU6 | 6IA2 | 6JQ5 | **6JXM** |
| **6MJ0** | **6N2V** | **6N5P** | 6OL3 | **6P2H** | 6PMO | 6QN3 | 6R47 | 6UFG | | | | |

[a] PDB IDs of the 35 RNAs in the training decoy set are marked in bold.

Table S2. Number of identified native structures by cgRNASP and other existing statistical potentials.[a]

| RNA data sets | Statistical potentials | | | | | | |
|---|---|---|---|---|---|---|---|
| | cgRNASP | rsRNASP | RNA3DCNN | ARES | DFIRE-RNA | 3dRNAscore | RASP |
| Test set_MD | 4/5 | 3/5 | 4 /5 | 0/5 | 3/5 | **5**/5 | 1/5 |
| Test set_NM | 11/15 | 13 /15 | **15**/15 | 5/15 | 12 /15 | 12/15 | 11/15 |
| Test set_PM | 15/22 | **16**/22 | 14/22 | 0/22 | 10 /22 | 2/22 | 2/22 |
| Test set_Puzzles | **18**/20 | 16/20 | 13/20 | 2/20 | 10/20 | 2/20 | 2/20 |
| Total[b] | **48**/62 | **48**/62 | 46/62 | 7/62 | 35/62 | 21/62 | 16/62 |

[a] Bold values mean that the best value obtained by these statistical potentials.

[b] Total includes all the test sets.

Table S3. Mean DI of lowest-energy structures including native ones by cgRNASP and other statistical potentials.[a]

| RNA data sets | Statistical potentials | | | | | | |
|---|---|---|---|---|---|---|---|
| | cgRNASP | rsRNASP | RNA3DCNN | ARES | DFIRE-RNA | 3dRNAscore | RASP |
| Test set_MD | 0.1 | 0.4 | 0.2 | 2.2 | 1.1 | **0.0** | 1.0 |
| Test set_NM | 0.6 | 0.2 | **0.0** | 1.3 | 0.4 | 0.4 | 0.5 |
| Test set_PM | 5.2 | **3.3** | 6.2 | 15.3 | 10.2 | 15.9 | 17.0 |
| Test set_Puzzles | **1.5** | 4.6 | 5.9 | 18.1 | 7.6 | 17.1 | 17.8 |
| Average on all[b] | **1.8** | 2.1 | 3.1 | 9.2 | 4.8 | 8.3 | 9.1 |
| Average on Puzzles+PM[c] | **3.3** | 3.9 | 6.1 | 16.7 | 8.9 | 16.5 | 17.4 |

[a] Bold values mean that the best ones obtained by the statistical potentials.

[b] Average value on the mean values of all test sets including MD, NM, PM, and Puzzles.

[c] Average value on the mean values of realistic test sets PM and Puzzles.

Table S4. Mean DI of lowest-energy structures excluding native ones by cgRNASP and other statistical potentials.[a]

| RNA data sets | Statistical potentials | | | | | | |
|---|---|---|---|---|---|---|---|
| | cgRNASP | rsRNASP | RNA3DCNN | ARES | DFIRE-RNA | 3dRNAscore | RASP |
| Test set_MD | **0.3** | 0.6 | 0.4 | 2.2 | 2.0 | **0.3** | 1.0 |
| Test set_NM | 1.8 | 1.6 | **1.5** | 1.8 | 1.6 | **1.5** | 1.6 |
| Test set_PM | 12.1 | **11.2** | 12.6 | 15.3 | 13.1 | 17.2 | 18.7 |
| Test set_Puzzles | **13.7** | 14.4 | 18.5 | 18.8 | 14.4 | 19.4 | 20.0 |
| Average on all[b] | 7.0 | **6.9** | 8.3 | 9.5 | 7.8 | 9.6 | 10.4 |
| Average on Puzzles+PM[c] | 12.9 | **12.8** | 15.6 | 17.1 | 13.8 | 18.3 | 19.4 |

[a] Bold values mean that the best ones obtained by the statistical potentials.

[b] Average value on the mean values of all test sets including MD, NM, PM, and Puzzles.

[c] Average value on the mean values of realistic test sets PM and Puzzles.

Table S5. Mean Pearson correlation coefficient by cgRNASP and other statistical potentials.[a]

| RNA data sets | Statistical potentials | | | | | | |
|---|---|---|---|---|---|---|---|
| | cgRNASP | rsRNASP | RNA3DCNN | ARES | DFIRE-RNA | 3dRNAscore | RASP |
| Test set_MD | 0.79 | 0.81 | 0.73 | 0.77 | **0.82** | 0.75 | 0.74 |
| Test set_NM | 0.83 | 0.89 | 0.88 | 0.88 | **0.91** | 0.90 | 0.87 |
| Test set_PM | 0.59 | **0.64** | 0.47 | 0.36 | 0.54 | 0.19 | 0.14 |
| Test set_Puzzles | **0.58** | 0.57 | 0.35 | 0.40 | 0.52 | 0.35 | 0.38 |
| Average on all[b] | 0.70 | **0.73** | 0.61 | 0.60 | 0.70 | 0.55 | 0.53 |
| Average on Puzzles+PM[c] | 0.59 | **0.61** | 0.41 | 0.38 | 0.53 | 0.27 | 0.26 |

[a] Bold values mean that the best ones obtained by the statistical potentials.

[b] Average value on the mean values of all test sets including MD, NM, PM, and Puzzles.

[c] Average value on the mean values of realistic test sets PM and Puzzles.

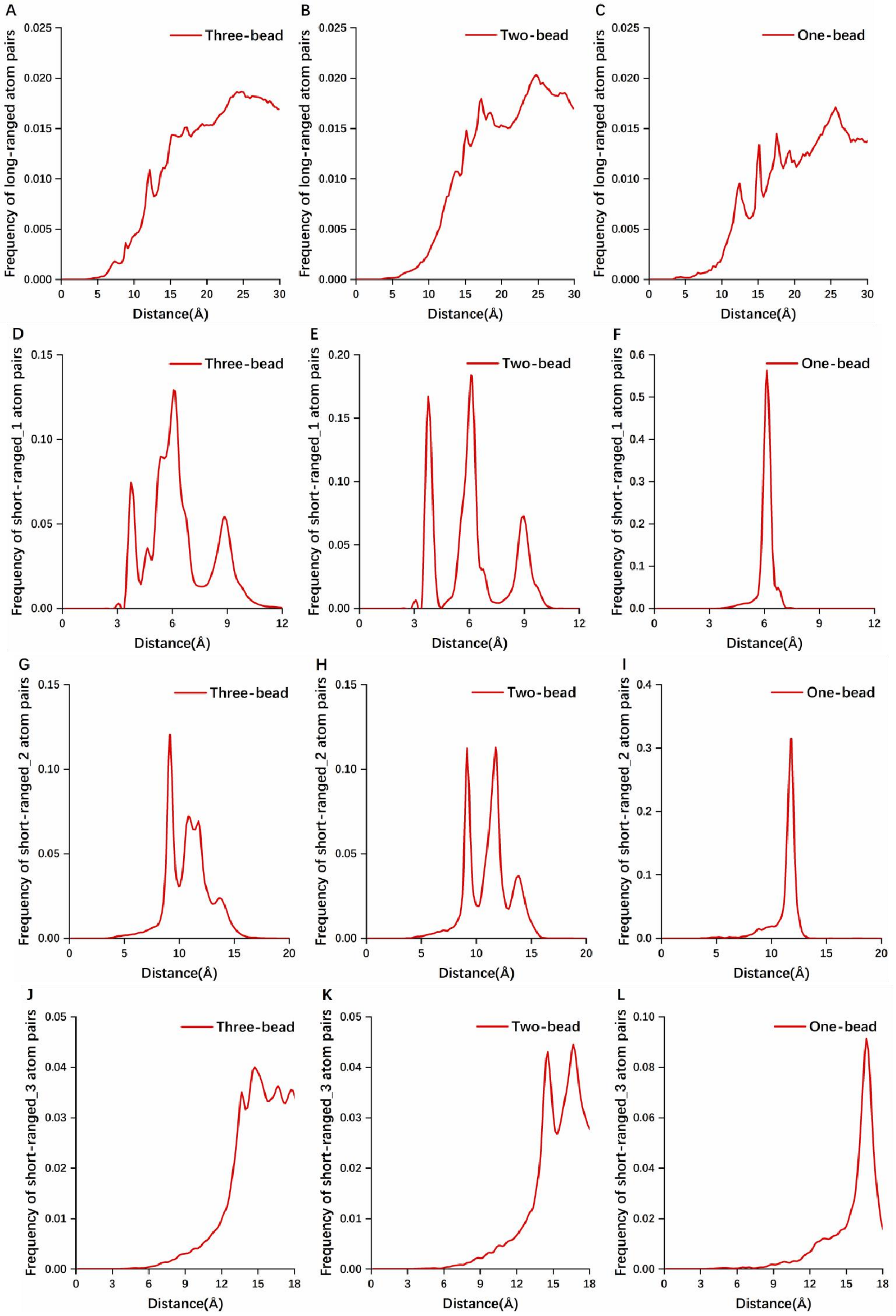


Figure S1. Distance distributions of long-ranged and short-ranged CG bead pairs from the statistics on the 191 native RNAs in the training native set for different CG representations used in cgRNASP potentials (cgRNASP, cgRNASP-PC, and cgRNASP-C); see the details in Section 2 in the Supplementary Material. (A-C) For residue separation of $k \geqslant 5$; (D-F) for residue separation of $k$=1; (G-I) for residue separation of $k$=2; and (J-L) for residue separation of $5>k\geqslant 3$.

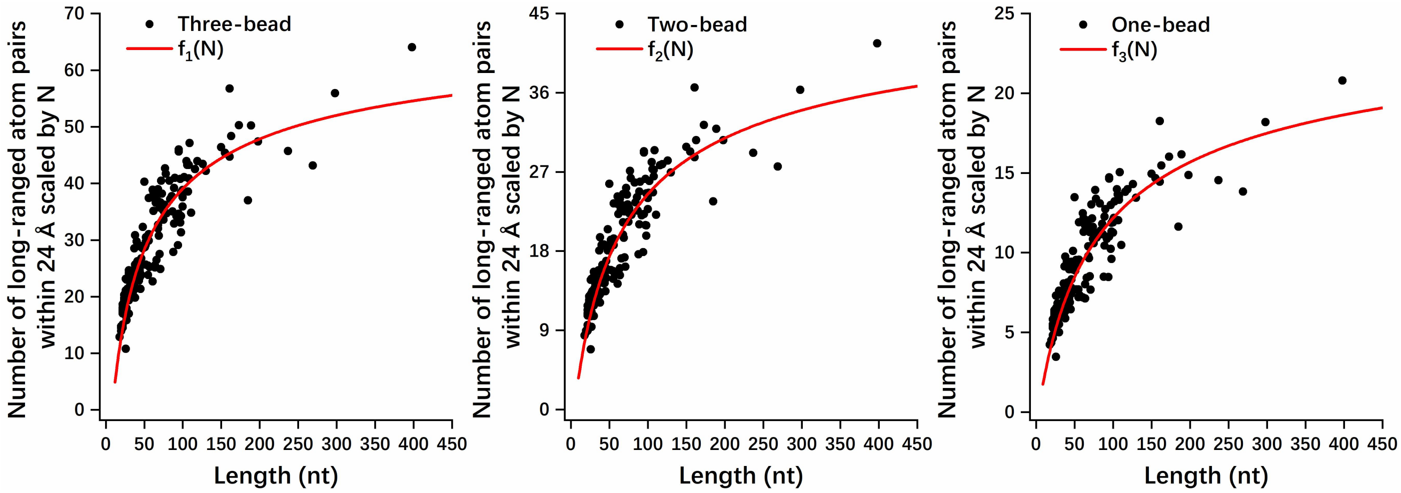


Figure S3. Relationships between the number of long-ranged CG bead pairs within 24 Å ($r_c$) per nucleotide and RNA length ($N$, in nt) from the statistics on the 191 native RNAs in the native training set; see the details in Section 2 in the Supplementary Material, and panels (A-C) are for different CG representations. Here, and $f_1(N)=\frac{-355}{(N+16)^{0.5}}+72$, $f_2(N)=\frac{-290}{(N+29)^{0.5}}+50$, and $f_3(N)=\frac{-175}{(N+39)^{0.5}}+27$ denote the fitted lines for panels (a), (b) and (c).

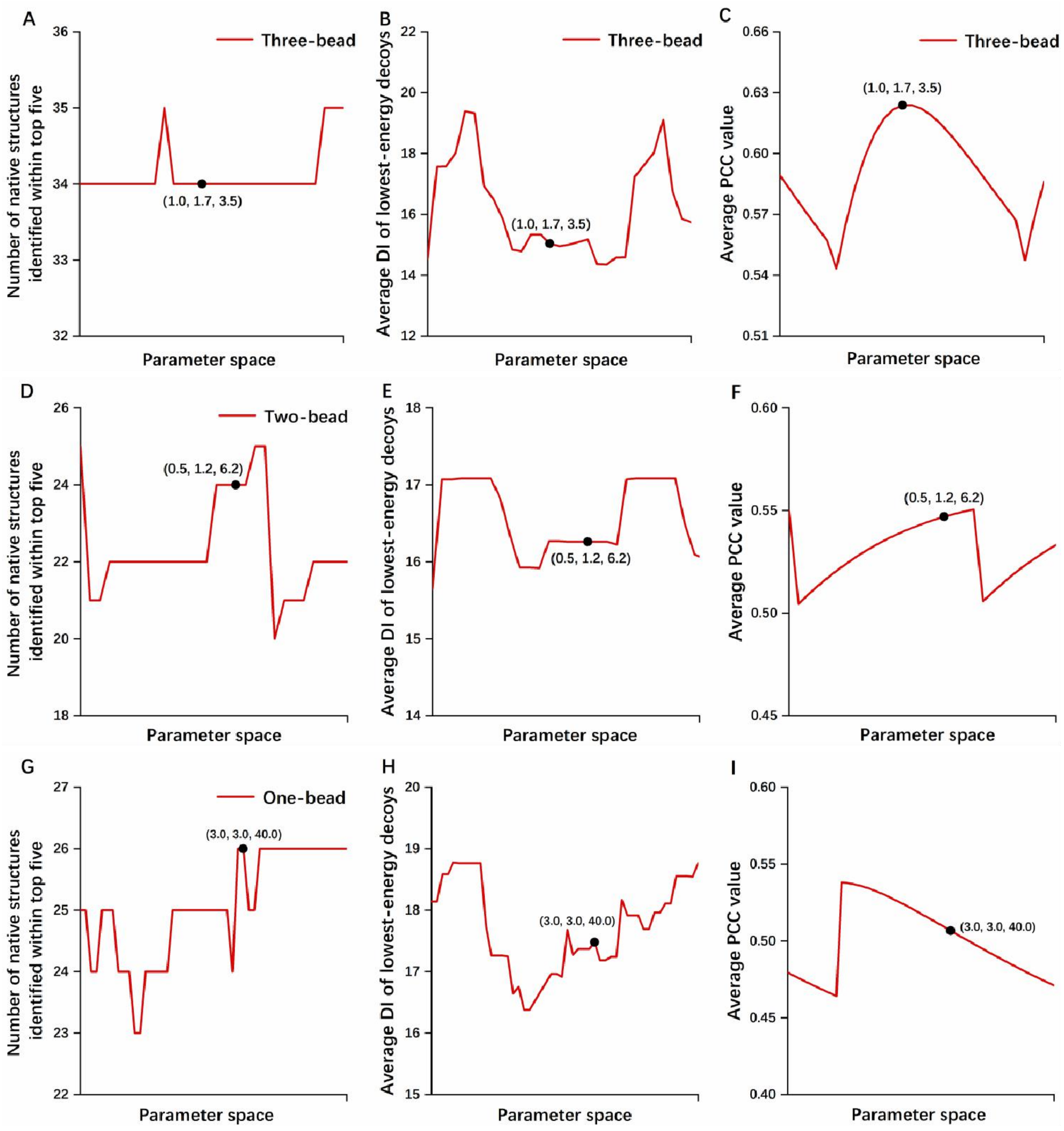


Figure S4. Illustration for the choice of the parameters (α, β, $w_0$) for different CG statistical potentials (cgRNASP, cgRNASP-PC, and cgRNASP-P) by three metrics (number of native structures identified within top five (lowest-energy) structures, average DI of lowest-energy decoy structures, and average PCC value on DI's) for the training decoy set, respectively. Please see the details in Section 2 in the Supplementary Material. (A-C) α=1.0, β=1.7 and $w_0$=3.5 for cgRNASP; (D-F) α=0.5, β=1.2, and $w_0$=6.2 for cgRNASP-PC; (G-I) α=3.0, β=3.0, and $w_0$=40.0 for cgRNASP-C.

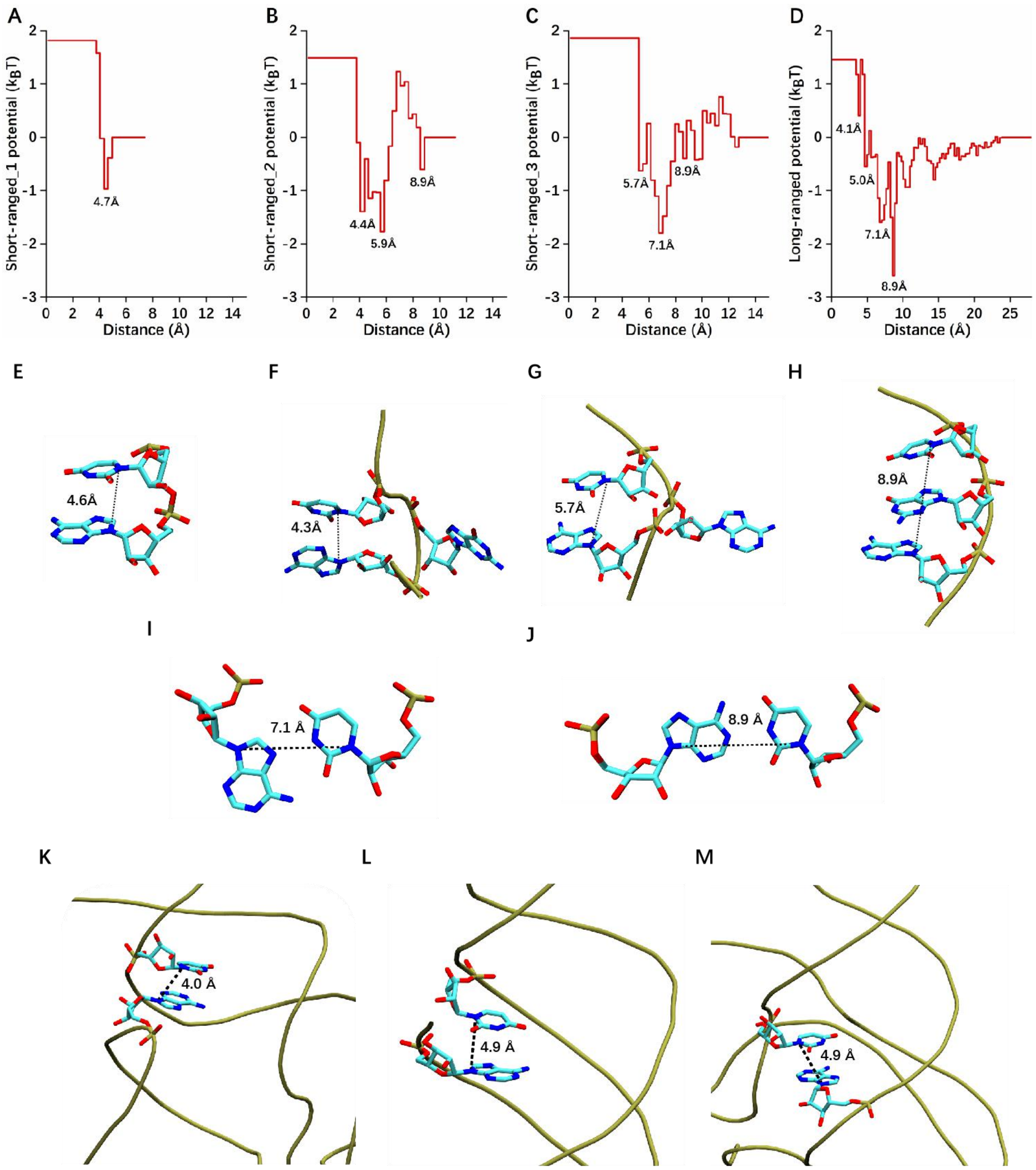


Figure S5 (A-C) Short-ranged and (D) long-ranged potentials between AN9 and UN1 in cgRNASP. In panels (A-D), short-ranged_1, short-ranged_2, and short-ranged_3 potentials denote those at residue separations of $k$=1, $k$=2, and $3 \le k \le 4$, respectively. (E-H) Representative distances between AN9 and UN1 for nearest-neighbor base stacking (E), next-nearest neighbor U-turns (F and G), and next-nearest neighbor base-stacking (H) captured in the short-ranged potentials. (I-M) Other representative distances for reverse Hoogsteen (~7.1 Å) and Watson-Crick base pairing (~8.9 Å) in the short-ranged_3 potential, as well as those for base stacking between adjacent branches (~4 Å), base stacking at triplex (~5 Å) and coaxial-stacking at junctions (~5 Å) in the long-ranged potential, which has been illustrated in Ref [1].
